## Supplementary Information for "Novel green fluorescent polyamines to analyze ATP13A2 and ATP13A3 activity in the mammalian polyamine transport system"

**This file contains:**

Supplementary synthetic methods and schemes

Supplementary $^{1}$H-NMR and $^{13}$C-NMR and HRMS data

Supplementary LC-chromatograms of final compounds including purity

### Synthetic methods

All starting materials and solvents were purchased from commercial vendors and used without further purification unless specified otherwise. Raney®-Nickel was purchased as a 50% slurry in water from Acros Organics (catalog number 395921000). Reactions were analysed by Thin Layer Chromatography (TLC) on pre-coated 0.20 mm thick ALUGRAM® TLC sheets with fluorescent indicator and by liquid chromatography mass spectrometry (LC MS) performed on a Prominence Ultra fast Liquid Chromatography system equipped with a 2x150 mm C18 analytical column (Waters X Bridge) coupled to a MS-2020 single quadrupole mass analyser (Shimadzu). A linear gradient of 5 to 90% acetonitrile in water (containing 0.1% formic acid) was used. Purity of compounds was assessed using 215 nm HPLC trace. The injection peak and peaks determined to be baseline variations that appear in all spectra were removed manually. HRMS spectra were acquired on a quadrupole orthogonal acceleration time of flight mass spectrometer (Synapt G2 HDMS, Waters, Milford, MA). Samples were infused at 3µL/min and spectra were obtained in positive ionization mode with a resolution of 15000 (FWHM) using leucine enkephalin as lock mass. Silica column chromatography was performed using 230-400 mesh silica (Kieselgel 60). NMR spectra were recorded in deuterated solvents as indicated and measured using a Bruker UltraShield 600 MHz NMR spectrometer. Chemical shifts are reported in ppm relative to the residual solvent peak and J-values are reported in Hertz.

The azide precursors were synthesized based on previously described procedures [1,2]. The symmetrical BODIPY probes were synthesized as HCl salts based on previously described procedures [1, 2].

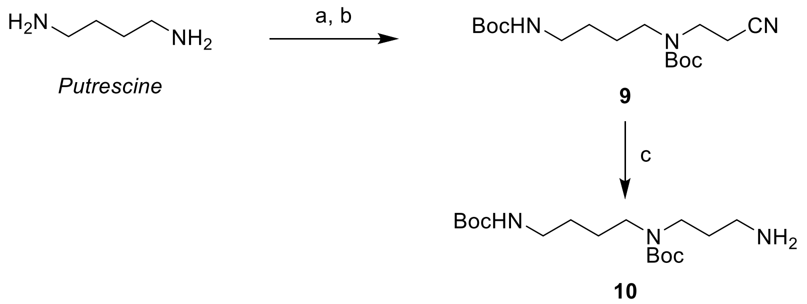


**Figure S5. Synthesis of putrescine precursors, reagents and conditions:** (a) Acrylonitrile, $Et_3N$, MeOH, 0 °C → rt. (b) $Boc_2O$, $Et_3N$, MeOH, 0 °C → rt. (c) Raney®-Nickel, NaOH, 2.3 bar of $H_2$, MeOH, rt.

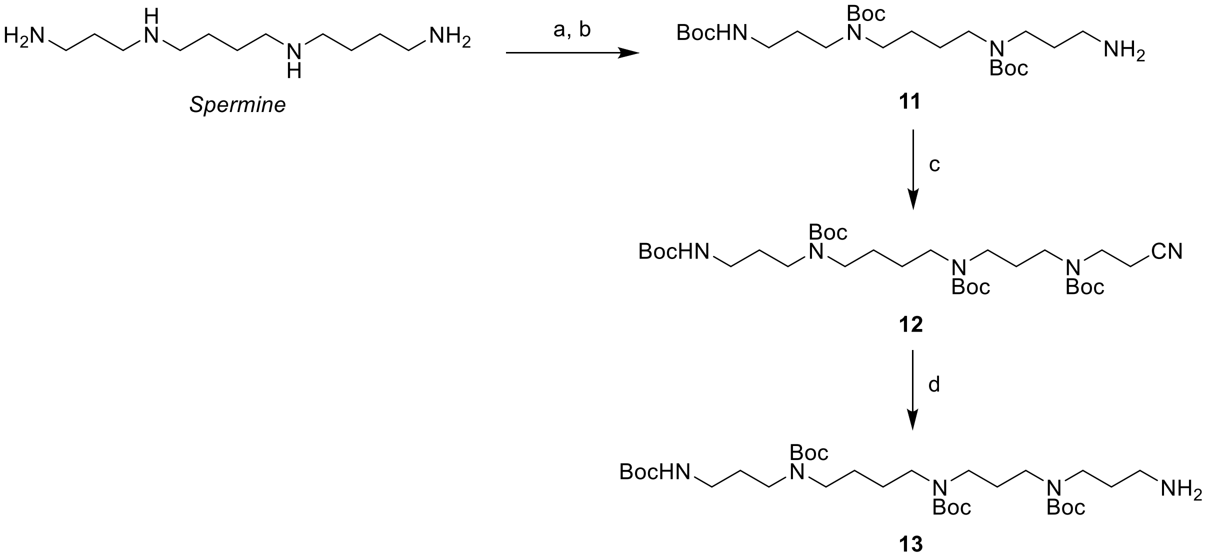


**Figure S6. Synthesis of spermidine and spermine precursors, reagents and conditions:** (a) EtTFA, MeOH, 0 °C → rt., then $Boc_2O$. (b) $NaOH_{(aq)}$, MeOH, rt. (c) acrylonitrile, 0 °C → rt., then $Boc_2O$. (d) Raney®-Nickel, ammonia, 3 bar $H_2$, MeOH, rt.

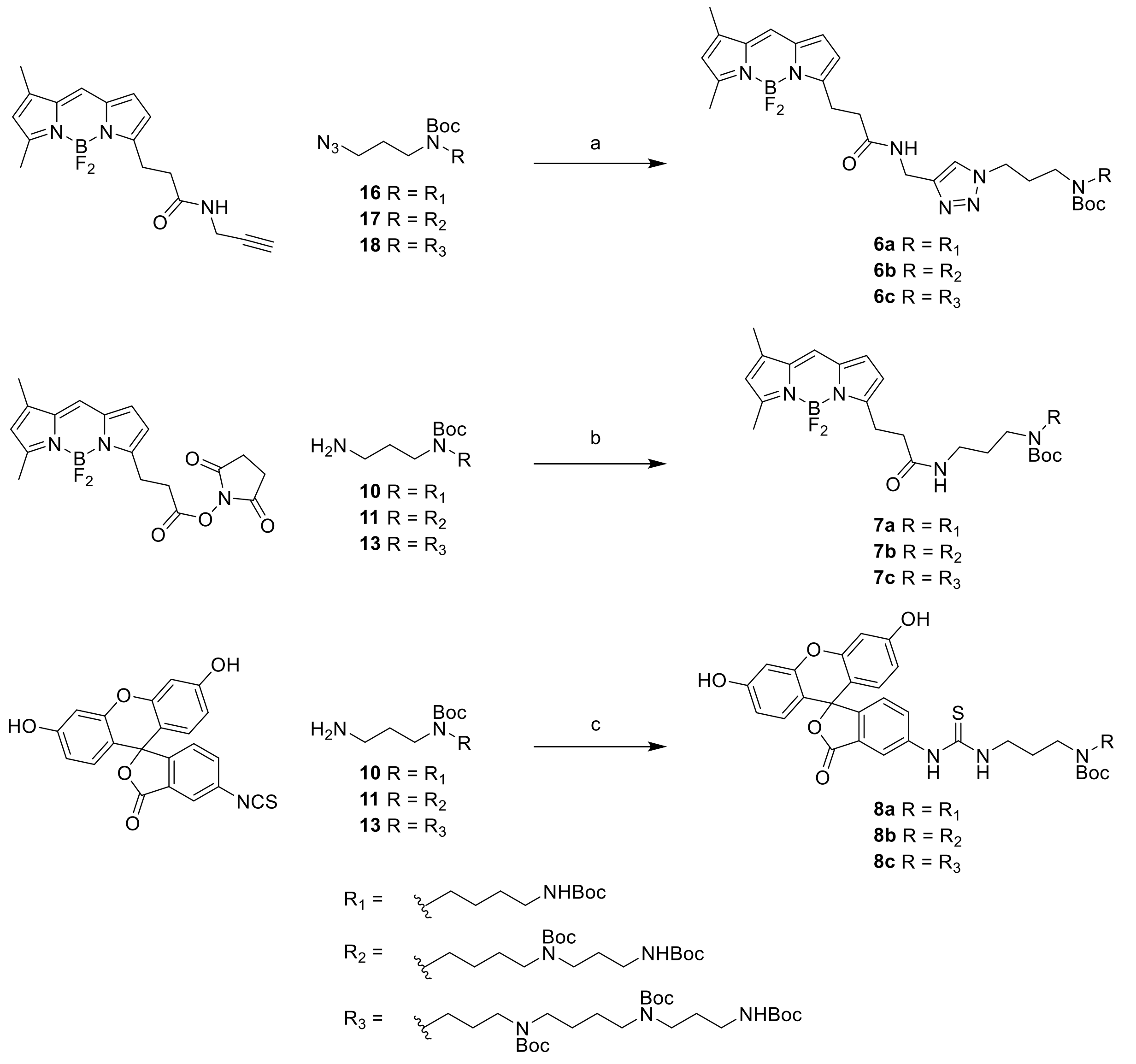


**Figure S7. Conjugation of the fluorophore to the relevant protected polyamine.** (a) CuBr, DIPEA, DCM, rt. (b) DIPEA, DCM, rt. (c) $Et_3N$, DCM, rt.

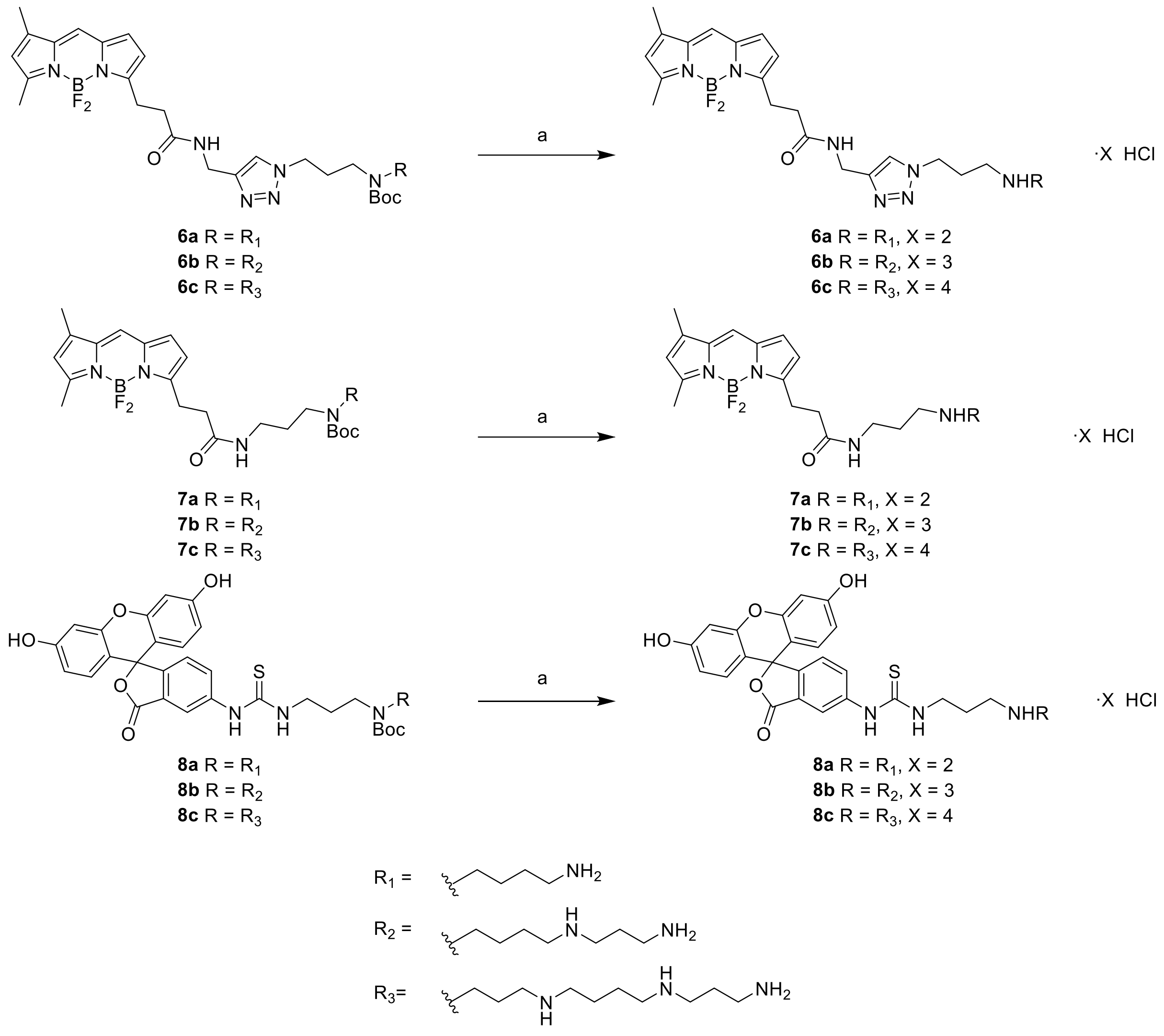


**Figure S8. Boc-deprotection of the relevant Boc-protected probe.** (a) 4 M HCl/dioxane, rt.

### General procedures

*General procedure A for synthesis of Boc protected BODIPY FL A probes*

The relevant Boc protected polyamine (1.2 eq) was dissolved in dichloromethane. Diisopropylethylamine (4.0 eq) was added followed by the BODIPY FL N hydroxysuccinimide ester (1.0 eq). The resulting solution was stirred overnight under argon atmosphere in the absence of light. The solvent was removed under a stream of nitrogen and the crude product was purified by silica chromatography (1/1 PE/EtOAc → PE/EtOAc 1/3).

*General procedure B for synthesis of Boc protected BODIPY FL T probes*

The relevant Boc protected polyamine (1.2 eq) was dissolved in dichloromethane. Diisopropylethylamine (4.0 eq) was added followed by copper bromide (0.5 eq). Subsequently, the relevant alkyne-BODIPY (1.0 eq) was added. The resulting suspension was stirred overnight under argon atmosphere in the absence of light. Following this, the mixture was washed three times using a saturated sodium bicarbonate solution. The solvent was then removed under a stream of nitrogen gas and the obtained crude product was purified using silica chromatography.

*General procedure C for synthesis of Boc protected FITC probes*

The relevant Boc protected polyamine (1.2 eq) was dissolved in dichloromethane. Triethylamine (4.0 eq) was added followed by FITC (1.0 eq). The resulting solution was stirred overnight under argon atmosphere in the absence of light. The solvent was removed under a stream of nitrogen and the crude product was purified by silica chromatography (5% MeOH in DCM + 1% AcOH).

*General procedure D for Boc deprotection towards the final polyamine probes*

1.0 mL 4 M HCl/dioxane was added to the relevant Boc protected polyamine probe (1.0 eq). The resulting solution was left to stand for 2.5 min per present Boc group generating colored precipitate. Following this, the solvent was removed in vacuo before dioxane was added. The resulting suspension was concentrated in vacuo. This was repeated two more times before the obtained solid was dissolved in water and lyophilised, yielding the final polyamine probe without further purification.

**_tert_-Butyl (4-((_tert_-butoxycarbonyl)amino)butyl)(2-cyanoethyl)carbamate (9)**

Putrescine (2.002 g, 1.0 eq) was dissolved in 18 mL of methanol after which the solution was purged *via* nitrogen gas bubbling for 20 minutes and left to stir under nitrogen atmosphere. The solution was cooled to 0 °C and acrylonitrile (1.5 mL, 1.0 eq) was added dropwise over the course of two hours after which the mixture was left to slowly warm to room temperature and further stirred for 24 hours. An additional 20 mL of methanol was added followed by triethylamine (20 mL, 6.3 eq). The mixture was cooled to 0 °C and Boc anhydride (15.039 g, 3.0 eq) was added in portions. The reaction was left to stir overnight at room temperature. The solvent was removed *in vacuo* and the resulting crude oil purified using silica chromatography (4/1 PE/EtOAc → PE/EtOAc 2/1). Pure fractions were pooled and concentrated *in vacuo* resulting in a colorless oil (2.07 g, yield = 26%).

ESI-MS: m/z calcd for $C_{17}H_{32}N_3O_4$ $[M+H]^+$: 342.24; found: 342.15

***tert*-Butyl (3-aminopropyl)(4-((*tert*-butoxycarbonyl)amino)butyl)carbamate (10)**

Nitrile **9** (318.3 mg, 1.0 eq) was added to a glass Parr® vessel and dissolved in 21 mL of 70% aqueous ethanol followed by the addition of 14 mL of a 1 M methanolic sodium hydroxide solution. Three scoops of Raney®-Nickel slurry (50% in water) were added and the formed suspension was put under an atmosphere of hydrogen at 2.3 bars and subsequently shaken overnight. The hydrogen atmosphere was removed and the suspension was filtered over a layer of Celite® 545. The filtrate was concentrated to a minimal volume *in vacuo* after which 75 mL of dichloromethane was added and washed twice using 75 mL of a 0.1 M aqueous sodium hydroxide and once using 75 mL of water. The organic phase was dried using magnesium sulfate, filtered, and concentrated resulting in a yellow oil (0.313 g, yield = 97%) which was used as such in subsequent reactions.

ESI-MS: m/z calcd for $C_{17}H_{36}N_3O_4$ $[M+H]^+$: 346.27; found: 346.90

***tert*-Butyl (4-((3-aminopropyl)(*tert*-butoxycarbonyl)amino)butyl)(3-((*tert*-butoxycarbonyl)amino) propyl)carbamate (11)**

Spermine (1.37 g, 1.3 eq) was dissolved in 40 mL methanol and put under argon atmosphere. Ethyl trifluoroacetate (0.643 mL, 1.0 eq) was dissolved in 20 mL methanol and added dropwise at 0 °C over the course of 30 minutes while stirring vigorously. After addition, the solution was left to warm to room temperature. After three hours of further stirring, Boc anhydride (8.890 g, 7.5 eq) was added in portions. The resulting solution was left to stir overnight under argon atmosphere. A 2 M aqueous sodium hydroxide solution was added dropwise resulting in the formation of a white precipitate. This suspension was further stirred overnight under argon atmosphere after which the mixture was concentrated to a minimal volume. Water was added to obtain a total volume of 150 mL which was extracted three times using 150 mL chloroform. The organic phases were combined, dried using magnesium sulfate, filtered, and concentrated *in vacuo*. The resulting crude material was purified using silica chromatography (2% MeOH in DCM → 5 % MeOH in DCM → 5 % MeOH in DCM + 1% $Et_3N$ → 10% MeOH in DCM + 1% $Et_3N$) resulting in a colorless oil (1.639 g, yield = 60%).

ESI-MS: m/z calcd for $C_{25}H_{51}N_4O_6$ $[M+H]^+$: 503.38; found: 503.35

***tert*-Butyl (4-((*tert*-butoxycarbonyl)(3-((*tert*-butoxycarbonyl)(2-cyanoethyl)amino)propyl)a mino)butyl)(3-((*tert*-butoxycarbonyl)amino)propyl)carbamate (12)**

Acrylonitrile (0.146 mL, 1.0 eq) was added dropwise to a solution of amine **11** (1.122 g, 1.0 eq) in 6 mL methanol, cooled to 0 °C. The resulting solution was stirred overnight under argon atmosphere after which triethylamine (1.24 mL, 4.0 eq) was added followed by the addition of Boc anhydride (1.43 g, 2.9 eq) in small portions. The solution was stirred overnight before being concentrated *in vacuo*. The crude mixture was purified using silica column chromatography (2/1 PE/EtOAC → 1/2 PE/EtOAc) resulting in a colorless oil (0.974, yield = 67%).

ESI-MS: m/z calcd for $C_{33}H_{62}N_5O_8$ $[M+H]^+$: 656.46; found: 656.45

***tert*-Butyl (4-((3-((3-aminopropyl)(*tert*-butoxycarbonyl)amino)propyl)(*tert*-butoxycarbonyl)amino)butyl)(3-((*tert*-butoxycarbonyl)amino)propyl)carbamate (13)**

Nitrile **12** (0.974 g, 1.0 eq) was dissolved in 10 mL of a 2 M methanolic ammonia solution followed by the addition of three scoops of a Raney®-Nickel slurry (50% in water). The resulting suspension was shaken overnight in a glass Parr® vessel under a hydrogen atmosphere of 3 bar. After removal of the hydrogen atmosphere solids were filtered off over Celite® 545. The filtrate was concentrated *in vacuo* and the obtained crude purified using silica chromatography (5% MeOH in DCM + 1% $Et_3N$). Clean fractions were pooled resulting in a colorless oil (0.616 g, yield = 63%).

ESI-MS: m/z calcd for $C_{33}H_{66}N_5O_8$ $[M+H]^+$: 660.49; found: 660.50

***tert*-Butyl (4-((*tert*-butoxycarbonyl)amino)butyl)(3-(4-((3-(5,5-difluoro-7,9-dimethyl-5*H*-4λ$^4$,5λ$^4$-dipyrrolo[1,2-c:2',1'-*f*][1,3,2]diazaborinin-3-yl)propanamido)methyl)-1*H*-1,2,3-triazol-1-yl)propyl)carbamate (6a)**

Prepared according to general procedure B using azide **16** (10.76 mg) as starting material. The crude was purified by silica chromatography (3% MeOH in DCM → 5% MeOH in DCM) resulting in a red solid (12.28 mg, yield = 61%).

$^1$H-NMR (600 MHz, DMSO-d6): δ = 8.42   8.37 (m, 1H), 7.88 (s, 1H), 7.64 (s, 1H), 7.03 (d, 1H, J = 3.76 Hz), 6.77 – 6.72 (m, 1H), 6.29 (d, 1H, J = 3.80 Hz), 6.26 (s, 1H), 4.29 – 4.22 (m, 4H), 3.13 – 3.01 (m, 7H), 2.89 – 2.82 (m, 2 H), 2.51 (s, 2H), 2.42 (s, 3H), 2.21 (s, 3H), 1.98 – 1.91 (m, 2H), 1.41 – 1.22 (m, 19H), 1.13 (t, 2H, J = 7.27) $^{13}$C-NMR (150 MHz, DMSO-d6)(rotamers): δ = 170.78, 159.20, 157.72, 156.85, 155.63, 155.15, 154.76, 154.48, 144.84, 144.13, 134.46, 132.96, 128.89, 125.33, 122.81, 120.30, 116.62, 78.52, 77.37, 47.22, 46.35, 46.02, 45.74, 45.70, 43.84, 43.73, 34.35, 34.23, 33.60, 29.03, 28.74, 28.26, 28.01, 26.88, 25.54, 25.12, 23.92, 23.29, 18.06, 16.72, 14.51, 10.99, 8.61.

ESI-MS: m/z calcd for $C_{34}H_{52}BF_2N_8O_5$ $[M+H]^+$: 701.41; found: 701.40

***tert*-Butyl (4-((*tert*-butoxycarbonyl)(3-((*tert*-butoxycarbonyl)amino)propyl)amino)butyl)(3-(4-((3-(5,5-difluoro-7,9-dimethyl-5*H*-4λ$^4$,5λ$^4$-dipyrrolo[1,2-*c*:2',1'-*f*][1,3,2]diazaborinin-3-yl)propanamido)methyl)-1*H*-1,2,3-triazol-1-yl)propyl)carbamate (6b)**

Prepared according to general procedure B using azide **17** (10.16 mg) as starting material. The crude was purified by silica chromatography (3% MeOH in DCM → 5% MeOH in DCM) resulting in a red solid (9.90 mg, yield = 65%).

$^1$H-NMR (600 MHz, DMSO-d6): δ = 8.36 (s, 1H), 7.88 (s, 1H), 7.60 (s, 1H), 7.00 (d, 1H, J = 3.82 Hz), 6.67 (br s, 1H), 6.26 (d, 1H, J = 3.90 Hz), 6.23 (s, 1H), 4.29 – 4.17 (m, 4H), 3.11 – 2.96 (m, 10H), 2.83 – 2.77 (m, 2H), 2.39 (s, 3H), 2.18 (s, 3H), 1.96 – 1.87 (m, 2H), 1.47 (br s, 2H), 1.37 – 1.19 (br, s 33H) $^{13}$C-NMR (150 MHz, DMSO-d6)(rotamers): δ = 170.81, 159.20, 157.66, 155.55, 154.65, 144.13, 134.44, 132.93, 128.86, 125.30, 123.13, 120.29, 116.59, 78.52, 78.29, 77.46, 47.23, 46.47, 45.94, 44.40, 44.18, 43.84, 37.58, 34.26, 33.62, 28.90, 28.22, 28.01, 27.97, 25.57, 25.08, 23.90, 14.48, 10.97.

ESI-MS: m/z calcd for $C_{42}H_{67}BF_2N_9O_7$ [M+H]$^+$: 858.52; found: 858.50

***tert*-Butyl (4-((*tert*-butoxycarbonyl)(3-((*tert*-butoxycarbonyl)amino)propyl)amino)butyl)(3-((*tert*-butoxycarbonyl)(3-(4-((3-(5,5-difluoro-7,9-dimethyl-5*H*-4λ$^4$,5λ$^4$-dipyrrolo[1,2-*c*:2',1'-*f*][1,3,2]diazaborinin-3-yl)propanamido)methyl)-1*H*-1,2,3-triazol-1-yl)propyl)carbamate (6c)**

Prepared according to general procedure B using azide **18** (10.12 mg) as starting material. The crude was purified by silica chromatography (3% MeOH in DCM → 5% MeOH in DCM) resulting in a red solid (4.12 mg, yield = 45%).

$^1$H-NMR (600 MHz, DMSO-d6): δ = 8.42 (s, 1H), 7.94 (s, 1H), 7.67 (s, 1H), 7.06 (d, 1H, J = 3.84), 6.72 (br s, 1H), 6.32 (d, 1H, J = 3.88), 6.29 (s, 1H), 4.34 – 4.25 (m, 3H), 3.18 – 3.01 (m, 16H), 2.90 – 2.84 (m, 2H), 2.45 (s, 3H), 2.25 (s, 3H), 2.03 – 1.94 (m, 2H), 1.62 (br s, 2H), 1.54 (br s, 2H), 1.42 – 1.31 (m, 41H) $^{13}$C-NMR (150 MHz, DMSO-d6)(rotamers): δ = 170.79, 159.20, 157.67, 155.55, 154.64, 154.55, 144.12, 134.44, 132.94, 128.87, 125.30, 123.10, 120.29, 116.59, 78.60, 78.30, 78.27, 77.45, 47.20, 46.36, 45.93, 45.77, 44.36, 44.14 ,43.82, 37.57, 34.26, 33.62, 28.88, 25.65, 25.11, 23.90, 14.48, 10.97, 8.62.

ESI-MS: m/z calcd for $C_{50}H_{82}BF_2N_{10}O_9$ [M+H]$^+$: 1015.63; found: 1015.65

***tert*-Butyl (4-((*tert*-butoxycarbonyl)amino)butyl)(3-(3-(5,5-difluoro-7,9-dimethyl-5*H*-4λ$^4$,5λ$^4$-dipyrrolo[1,2-*c*:2',1'-*f*][1,3,2]diazaborinin-3-yl)propanamido)propyl)carbamate (7a)**

Prepared according to general procedure A using amine **10** (7.23 mg) as starting material resulting in a red solid (9.2 mg, yield = 80%).

$^{1}H$-NMR (600 MHz, $CDCl_3$): δ = 7.08 (s, 1H), 6.87 (d, 1H, *J* = 3.2 Hz), 6.29 (d, 1H, J = 3.8 Hz), 6.10 (s, 1H), 3.32 – 3.26 (m, 2H), 3.19 (q, 3H, J = 6.2), 3.17 – 3.08 (m, 6H), 2.63 (t, 2H, J = 7.47), 2.56 (s, 3H), 2.25 (s, 3H), 1.62 (quint, 2H, J = 6.4), 1.55 – 1.47 (m, 1H), 1.43 (s, 19H), 1.36 – 1.30 (m, 1H) $^{13}C$-NMR (150 MHz, $CDCl_3$)(rotamers): δ = 171.84, 156.14, 143.69, 133.50, 128.40, 123.88, 120.39, 117.47, 79.76, 79.28, 46.82, 45.78, 44.86, 43.78, 40.27, 37.23, 36.16, 35.96, 32.06, 31.57, 30.33, 29.84, 29.50, 29.40, 28.56, 27.98, 27.54, 25.90, 24.90, 22.83, 15.05, 14.26, 11.44, 8.65.

ESI-MS: m/z calcd for $C_{31}H_{48}BF_2N_5O_5$ $[M+H]^+$: 620.38; found: 620.40

***tert*-Butyl (4-((*tert*-butoxycarbonyl)(3-(3-(5,5-difluoro-7,9-dimethyl-5*H*-4λ$^4$,5λ$^4$-dipyrrolo[1,2-*c*:2',1'-*f*][1,3,2]diazaborinin-3-yl)propanamido)propyl)amino)butyl)(3-((*tert*-butoxycarbonyl)amino)propyl)carbamate (7b)**

Prepared according to general procedure A using amine **11** (7.08 mg) as starting material resulting in a red solid (10.68 mg, yield = 76%).

$^{1}H$-NMR (600 MHz, $CDCl_3$): δ = 7.07 (s, 1H), 6.87 (d, 1H, J = 2.3 Hz), 6.29 (d, 1H, J = 3.6 Hz), 6.10 (s, 1H), 3.34 – 3.02 (m, 14H), 2.62 (t, 2H, J = 7.44), 2.55 (s, 3H), 2.24 (s, 1H), 1.67 – 1.58 (m, 4H), 1.51 – 1.36 (m, 30H), 1.36 – 1.29 (m, 1H) $^{13}C$-NMR (150 MHz, $CDCl_3$)(rotamers): δ = 171.85, 156.27, 155.57, 143.62, 135.17, 133.48, 128.38, 123.86, 120.38, 117.42, 79.75, 46.95, 46.35, 45.76, 44.83, 44.30, 43.76, 37.81, 37.51, 36.11, 35.94, 32.06, 31.57, 30.32, 29.83, 29.49, 29.40, 29.02, 28.58, 27.96, 26.21, 25.54, 24.89, 22.82, 15.05, 14.25, 11.44, 8.64.

ESI-MS: m/z calcd for $C_{39}H_{64}BF_2N_6O_7$ $[M+H]^+$: 777.49; found: 777.50

***tert*-Butyl (4-((*tert*-butoxycarbonyl)(3-((*tert*-butoxycarbonyl)(3-(3-(5,5-difluoro-7,9-dimethyl-5*H*-4λ$^4$,5λ$^4$-dipyrrolo[1,2-c:2',1'-f][1,3,2]diazaborinin-3-yl)propanamido)propyl)amino)propyl)amino)butyl)(3-((*tert*-butoxycarbonyl)amino)propyl)carbamate (7c)**

Prepared according to general procedure A using amine **18** (7.21 mg) as starting material resulting in a red solid (17.09 mg, yield = 99%).

$^{1}H$-NMR (600 MHz, $CDCl_3$): δ = 7.07 (s, 1H), 6.87 (d, 1H, J = 3.5 Hz), 6.28 (d, 1H, J = 4.00 Hz), 6.10 (s, 1H), 3.32 – 3.03 (m, 18H), 2.61 (t, 2H, J = 7.58 Hz), 2.54 (s, 3H), 2.24 (s, 3H), 1.75 – 1.59 (m, 6H), 1.50 – 1.37 (m, 40H) $^{13}C$-NMR (150 MHz, $CDCl_3$)(rotamers): δ = 171.80, 170.82, 160.02, 158.24, 157.64, 156.23, 155.57, 143.63, 137.26, 135.15, 133.48, 131.04, 129.61, 128.95, 128.39, 124.58, 124.09, 123.87, 120.37, 119.21, 117.42, 99.49, 79.82, 79.62, 79.51, 79.23, 79.01, 66.32, 58.00, 54.50, 46.95, 46.64, 46.48, 44.96, 44.30, 43.72, 39.49, 39.19, 39.03, 38.97, 37.80, 37.47, 37.22, 37.15, 36.05, 35.94, 32.05, 31.56, 28.57, 28.10, 27.89, 26.13, 25.66, 24.99, 24.87, 22.81, 17.75, 17.56, 17.37, 15.04, 14.24, 11.42, 10.15, 8.10.

ESI-MS: m/z calcd for $C_{47}H_{79}BF_2N_7O_9$ [M+H]$^+$: 934.60; found: 934.60

***tert*-Butyl (4-((*tert*-butoxycarbonyl)amino)butyl)(3-(3-(3',6'-dihydroxy-3-oxo-3*H*-spiro[isobenzofuran-1,9'-xanthen]-5-yl)thioureido)propyl)carbamate (8a)**

Prepared according to general procedure C using amine **14** (7.17 mg) as starting material resulting in a yellow solid (17.09 mg, yield = 99%).

$^1$H-NMR (600 MHz, $CD_3OD$): δ = 8.12 (s, 1H), 7.77 (d, 1H, J = 7.90 Hz), 7.18 (d, 1H, J = 8.18 Hz), 6.82 – 6.65 (m, 4H), 6.59 (d, 2H, J = 7.50 Hz), 3.63 (s, 2H), 3.23 (t, 2H, J = 7.32 Hz), 3.05 (t, 2H, J = 6.87), 1.98 – 1.81 (m, 2H), 1.62 – 1.53 (m, 2H), 1.51 – 1.37 (m, 23H) $^{13}$C-NMR (150 MHz, $CD_3OD$)(rotamers): δ = 210.11, 182.96, 182.61, 170.76, 162.16, 158.58, 157.83, 157.39, 154.75, 142.49, 142.20, 131.78, 130.68, 130.19, 129.37, 126.29, 120.71, 114.22, 112.04, 103.49, 81.08, 79.87, 48.09, 47.95, 46.23, 45.18, 43.50, 42.72, 42.36, 40.98, 30.76, 30.67, 29.35, 28.81, 28.79, 28.57, 28.37, 27.02, 26.52, 24.22.

ESI-MS: m/z calcd for $C_{38}H_{47}N_4O_9S$ [M+H]$^+$: 735.31; found: 735.35

***tert*-Butyl (4-((*tert*-butoxycarbonyl)(3-(3-(3',6'-dihydroxy-3-oxo-3*H*-spiro[isobenzofuran-1,9'-xanthen]-5-yl)thioureido)propyl)amino)butyl)(3-((*tert*-butoxycarbonyl)amino)propyl)carbamate (8b)**

Prepared according to general procedure C using amine **11** (7.43 mg) as starting material resulting in a yellow solid (17.09 mg, yield = 99%).

$^1$H-NMR (600 MHz, $CD_3OD$): δ = 8.09 (s, 1H), 7.74 (d, 1H, J = 7.71 Hz), 7.16 (d, 1H, J = 8.21 Hz), 6.74 – 6.65 (m, 4H), 6.54 (dd, 1H, J = 8.62 Hz, 1.94 Hz), 3.62 (s, 2H), 3.27 – 3.18 (m, 6H), 3.02 (t, J = 6.82 Hz), 1.95 – 1.80 (m, 2H), 1.68 (s, 2H), 1.58 – 1.37 (m, 33H) $^{13}$C-NMR (150 MHz, $CD_3OD$)(rotamers): δ = 210.09, 182.94, 182.61, 171.12, 161.74, 158.43, 157.77, 157.38, 154.30, 149.56, 142.30, 142.06, 131.94, 130.39, 129.45, 125.96, 120.30, 113.81, 111.58, 103.53, 81.07, 80.99, 79.97, 48.28, 47.91, 47.78, 46.26, 46.11, 45.66, 45.29, 43.51, 42.72, 38.93, 38.85, 30.77, 30.69, 30.26, 29.47, 28.82, 27.27, 27.14, 26.67, 26.53.

ESI-MS: m/z calcd for $C_{46}H_{62}N_5O_{11}S$ [M+H]$^+$: 892.42; found: 892.45

***tert*-Butyl (4-((*tert*-butoxycarbonyl)(3-((*tert*-butoxycarbonyl)(3-(3-(3',6'-dihydroxy-3-oxo-3*H*-spiro[isobenzofuran-1,9'-xanthen]-5-yl)thioureido)propyl)amino)propyl)amino)butyl)(3-((*tert* butoxycarbonyl)amino)propyl)carbamate (8c)**

Prepared according to general procedure C using amine **18** (7.18 mg) as starting material resulting in a yellow solid (19.08 mg, yield = 99%).

$^{1}$H-NMR (600 MHz, CD3OD): δ = 8.08 (s, 1H), 7.74 (d, 1H, J = 7.84 Hz), 7.16 (d, 1H, J = 8.22), 6.75 – 6.64 (m, 4H), 6.54 (dd, 2H, J = 8.70 Hz, 2.10 Hz), 3.63 (s, 2H), 3.27 – 3.12 (m, 12H), 3.02 (t, 2H, J = 6.69 Hz), 1.98 – 1.83 (m, 2H), 1.80 (q, 2H, J = 14.04 Hz), 1.67 (s, 2H), 1.55 – 1.38 (m, 40H) $^{13}$C-NMR (150 MHz, $CD_3OD$)(rotamers): δ = 210.11, 182.89, 182.59, 171.18, 169.33, 169.30, 162.33, 158.41, 157.30, 154.51, 148.80, 142.24, 142.06, 133.57, 132.41, 132.36, 131.66, 130.49, 129.87, 126.18, 120.44, 114.16, 111.80, 103.56, 81.17, 80.96, 79.95, 69.53, 48.26, 47.87, 47.75, 46.24, 46.05, 45.62, 45.24, 43.47, 42.74, 38.92, 38.83, 38.41, 34.80, 33.30, 33.06, 32.41, 30.75, 30.68, 30.61, 30.46, 30.29, 30.22, 30.13, 29.47, 28.84, 28.81, 27.54, 27.23, 27.05, 26.63, 23.72, 23.67, 20.97, 14.73, 14.44, 14.41, 9.20.

ESI-MS: m/z calcd for $C_{54}H_{77}N_6O_{13}S$ $[M+H]^+$: 1049.53; found: 1049.55

***N*-((1-(3-((4-Aminobutyl)amino)propyl)-1*H*-1,2,3-triazol-4-yl)methyl)-3-(5,5-difluoro-7,9-dimethyl-5*H*-4λ$^4$,5λ$^4$-dipyrrolo[1,2-*c*:2',1'-*f*][1,3,2]diazaborinin-3-yl)propenamide dihydrochloride (2a)**

Prepared according to general procedure D using Boc-protected probe **6a** (5.82 mg) as starting material. Obtained without purification as a red solid (4.65 mg, yield = 98%).

HRMS: m/z calcd for $C_{24}H_{36}BF_2N_8O$ $[M-H2Cl]^+$: 501.3073; found: 501.3074

***N*-((1-(3-((4-((3-Aminopropyl)amino)butyl)amino)propyl)-1*H*-1,2,3-triazol-4-yl)methyl)-3-(5,5-difluoro-7,9-dimethyl-5*H*-4λ$^4$,5λ$^4$-dipyrrolo[1,2-c:2',1'-f][1,3,2]diazaborinin-3-yl)propenamide trihydrochloride (2b)**

Prepared according to general procedure D using Boc-protected probe **6b** (7.06 mg) as starting material. Obtained without purification as a red solid (5.15 mg, yield = 94%).

HRMS: m/z calcd for $C_{27}H_{43}BF_2N_9O$ $[M-2H3Cl]^+$: 558.3651; found: 558.3652

***N*-((1-(3-((3-((4-((3-Aminopropyl)amino)butyl)amino)propyl)amino)propyl)-1*H*-1,2,3-triazol-4-yl)methyl)-3-(5,5-difluoro-7,9-dimethyl-5*H*-4λ$^4$,5λ$^4$-dipyrrolo[1,2-c:2',1'-f][1,3,2]diazaborinin-3-yl)propenamide tetrahydrochloride (2c)**

Prepared according to general procedure D using Boc-protected probe **6c** (5.29 mg) as starting material. Obtained without purification as a red solid (3.2 mg, yield = 81%).

HRMS: m/z calcd for $C_{30}H_{50}BF_2N_{10}O$ $[M-3H4Cl]^+$:615.4230; found: 615.4225

***N*-(3-((4-Aminobutyl)amino)propyl)-3-(5,5-difluoro-7,9-dimethyl-5*H*-4λ$^4$,5λ$^4$-dipyrrolo[1,2-*c*:2',1'-*f*][1,3,2]diazaborinin-3-yl)propenamide dihydrochloride (3a)**

Prepared according to general procedure D using Boc-protected probe **7a** (3.54 mg) as starting material. Obtained without purification as a red solid (3.84 mg, yield = 99%).

HRMS: m/z calcd for $C_{21}H_{33}BF_2N_5O$ [M-H2Cl]$^+$: 420.2746; found: 420.2747

***N*-(3-((4-((3-Aminopropyl)amino)butyl)amino)propyl)-3-(5,5-difluoro-7,9-dimethyl-5*H*-4λ$^4$,5λ$^4$-dipyrrolo[1,2-*c*:2',1'-*f*][1,3,2]diazaborinin-3-yl)propenamide trihydrochloride (3b)**

Prepared according to general procedure D using Boc-protected probe **7b** (3.22 mg) as starting material. Obtained without purification as a red solid (2.41 mg, yield = 99%).

HRMS: m/z calcd for $C_{24}H_{40}BF_2N_6O$ [M-2H3Cl]$^+$: 477.3324; found: 477.3322

***N*-(3-((3-((4-((3-Aminopropyl)amino)butyl)amino)propyl)amino)propyl)-3-(5,5-difluoro-7,9-dimethyl-5*H*-4λ$^4$,5λ$^4$-dipyrrolo[1,2-*c*:2',1'-*f*][1,3,2]diazaborinin-3-yl)propenamide tetrahydrochloride (3c)**

Prepared according to general procedure D using Boc-protected probe **7c** (4.01 mg) as starting material. Obtained without purification as a red solid (2.39 mg, yield = 97%).

HRMS: m/z calcd for $C_{27}H_{47}BF_2N_7O$ [M-3H4Cl]$^+$: 534.3903; found: 534.3909

**1-(3-((4-Aminobutyl)amino)propyl)-3-(3',6'-dihydroxy-3-oxo-3*H*-spiro[isobenzofuran-1,9'-xanthen]-5-yl)thiourea dihydrochloride (4a)**

Prepared according to general procedure D using Boc-protected probe **8a** (3.61 mg) as starting material. Obtained without purification as a yellow solid (2.92 mg, yield = 98%).

HRMS: m/z calcd for $C_{28}H_{31}N_4O_5S$ [M-H2Cl]$^+$: 535.2015; found: 535.2012

**1-(3-((4-((3-Aminopropyl)amino)butyl)amino)propyl)-3-(3',6'-dihydroxy-3-oxo-3*H*-spiro[isobenzofuran-1,9'-xanthen]-5-yl)thiourea trihydrochloride (4b)**

Prepared according to general procedure D using Boc-protected probe **8b** (4.80 mg) as starting material. Obtained without purification as a yellow solid (3.71 mg, yield = 98%).

HRMS: m/z calcd for $C_{31}H_{38}N_5O_5S$ [M-2H3Cl]$^+$: 592.2593; found: 592.2598

**1-(3-((3-((4-((3-Aminopropyl)amino)butyl)amino)propyl)amino)propyl)-3-(3',6'-dihydroxy-3-oxo-3*H*-spiro[isobenzofuran-1,9'-xanthen]-5-yl)thiourea tetrahydrochloride (4c)**

Prepared according to general procedure D using Boc-protected probe **8c** (4.95 mg) as starting material. Obtained without purification as a yellow solid (3.6 mg, yield = 96%).

HRMS: m/z calcd for $C_{34}H_{45}N_6O_5S$ [M-3H4Cl]+: 649.3172; found: 649.3174

### $^{1}$H-NMR and $^{13}$C-NMR data

Boc-protected BODIPY-FL-T **6a**

#### $^{1}$H-NMR

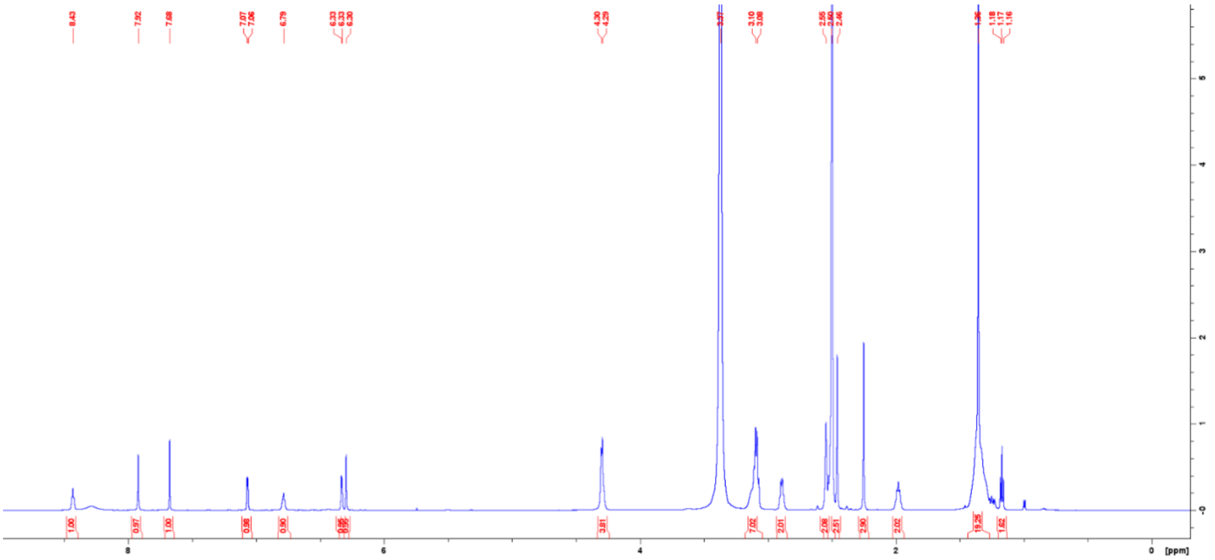

#### $^{13}$C-NMR

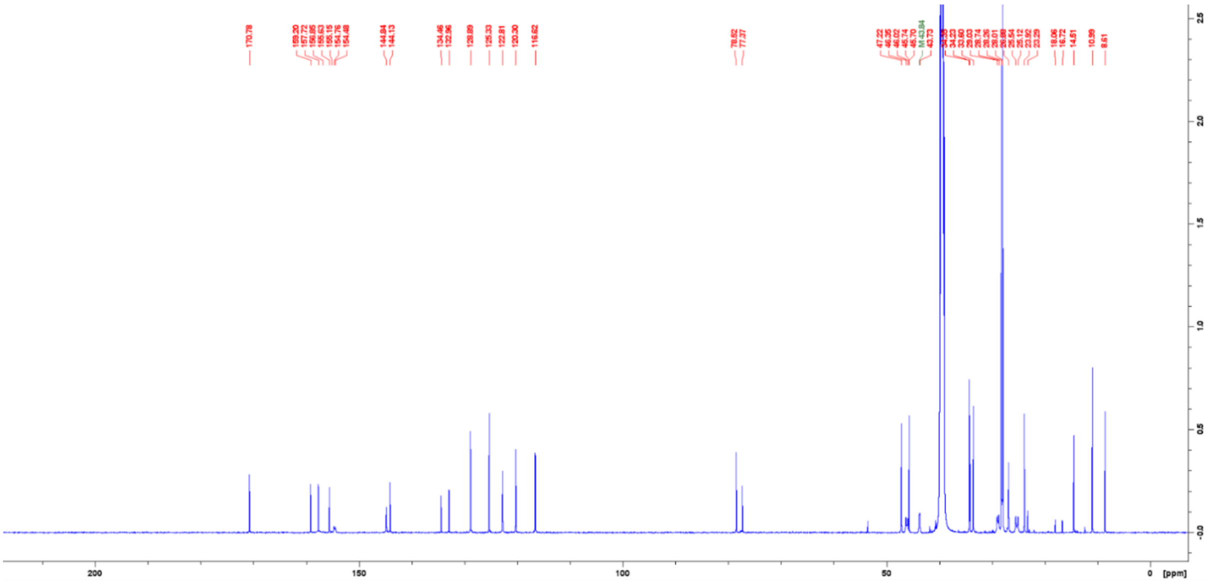

Boc-protected BODIPY-FL-T **6b**

### $^{1}$H-NMR

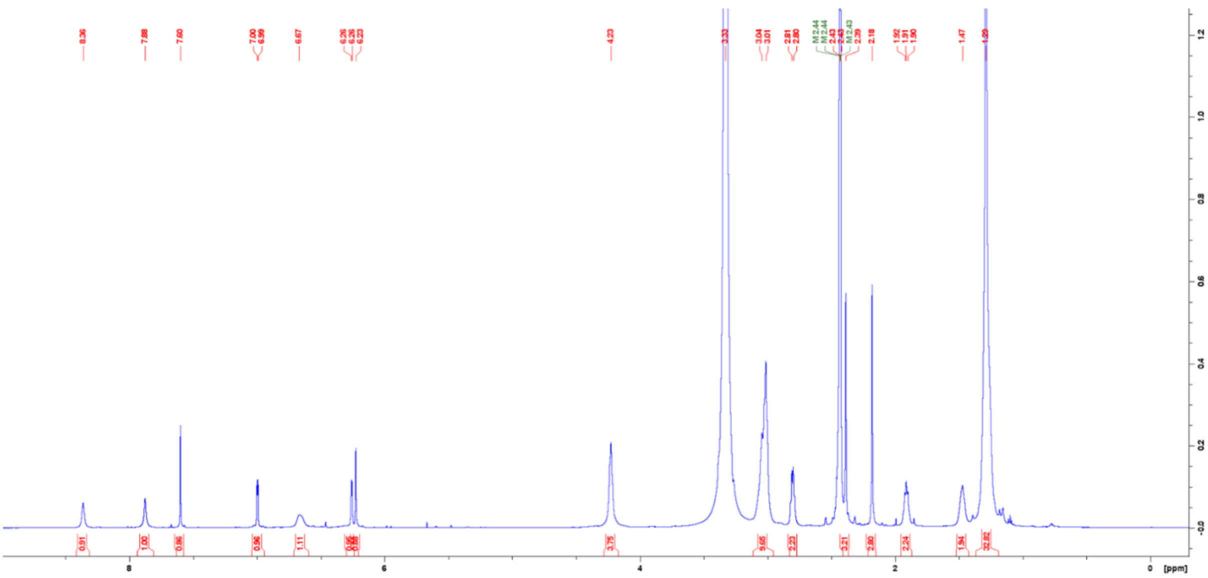


### $^{13}$C-NMR

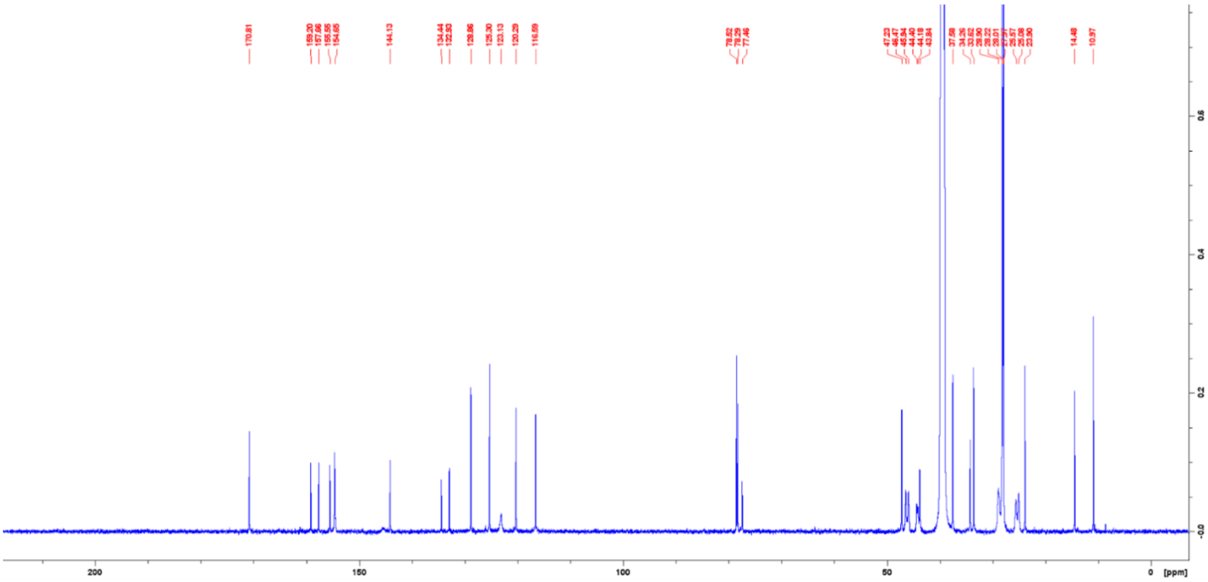

Boc-protected BODIPY-FL-T **6c**

**$^{1}$H-NMR**

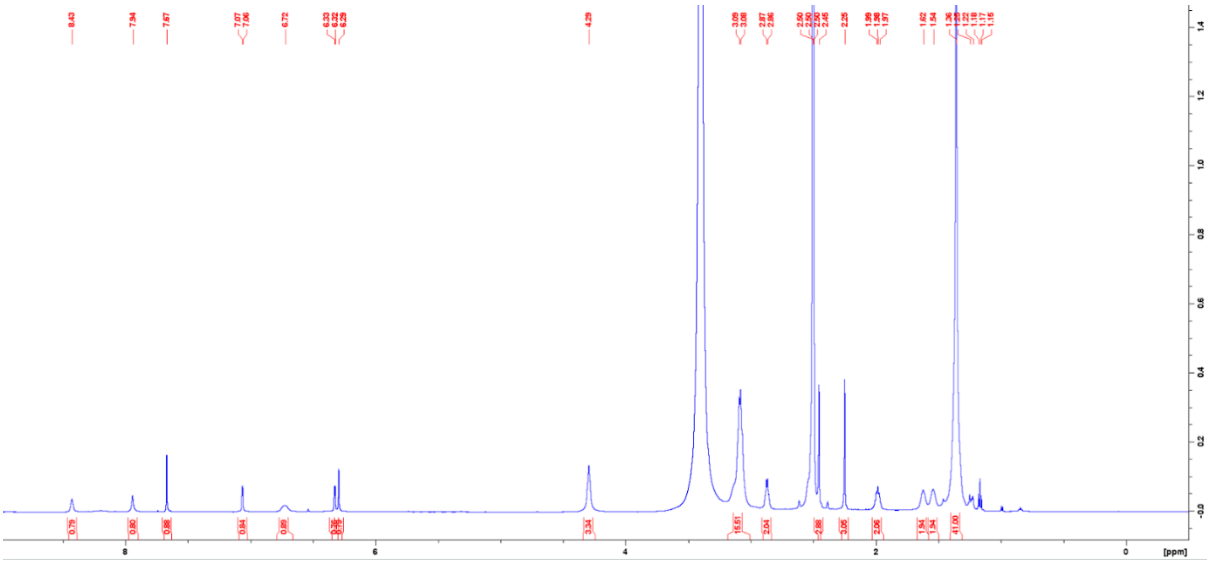

**$^{13}$C-NMR**

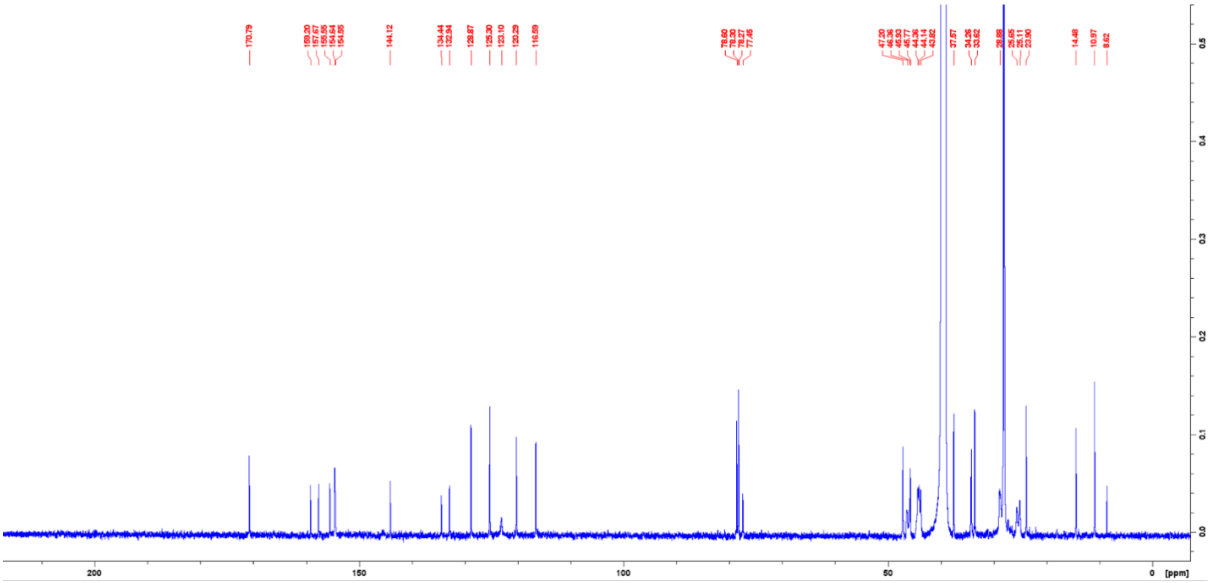

Boc-protected BODIPY-FL-A **7a**

**$^{1}$H-NMR**

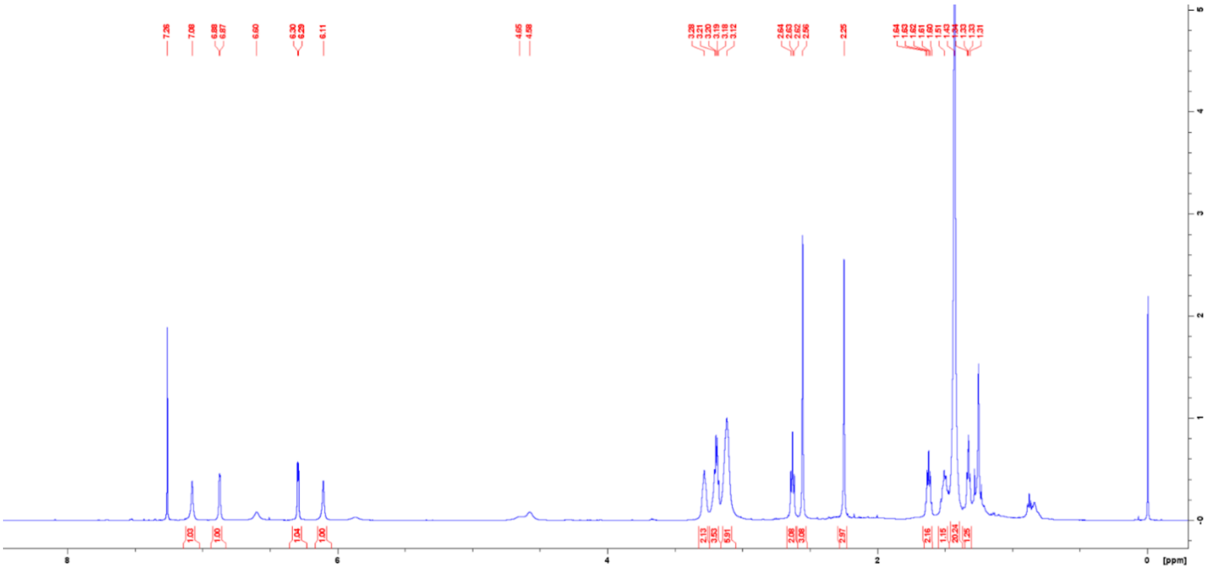

**$^{13}$C-NMR**

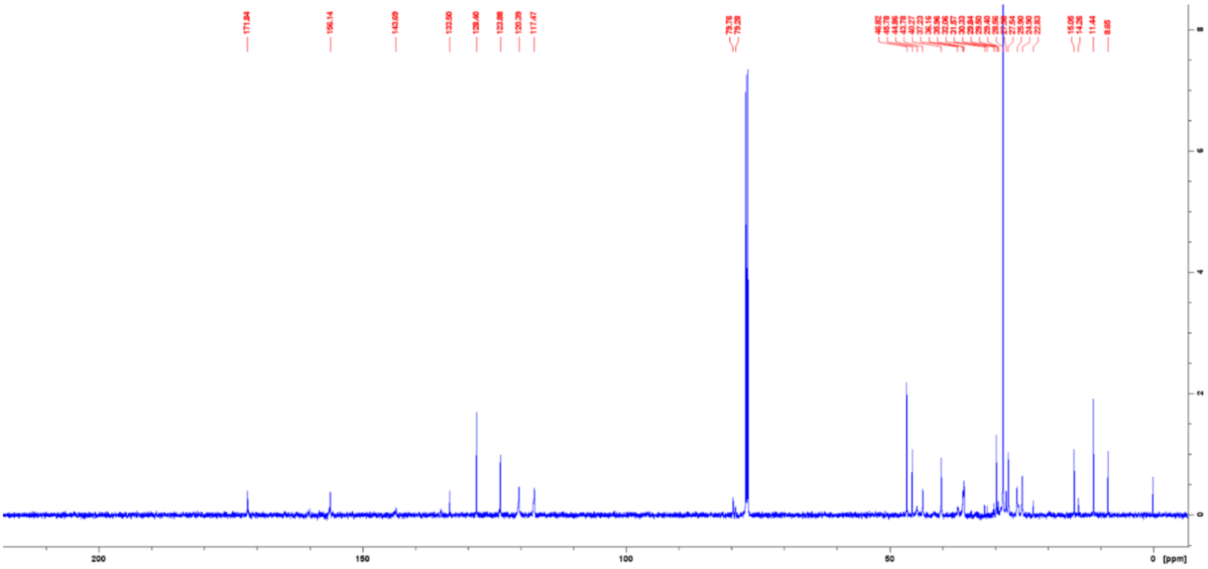

Boc-protected BODIPY-FL-A **7b**

**[1]H-NMR**

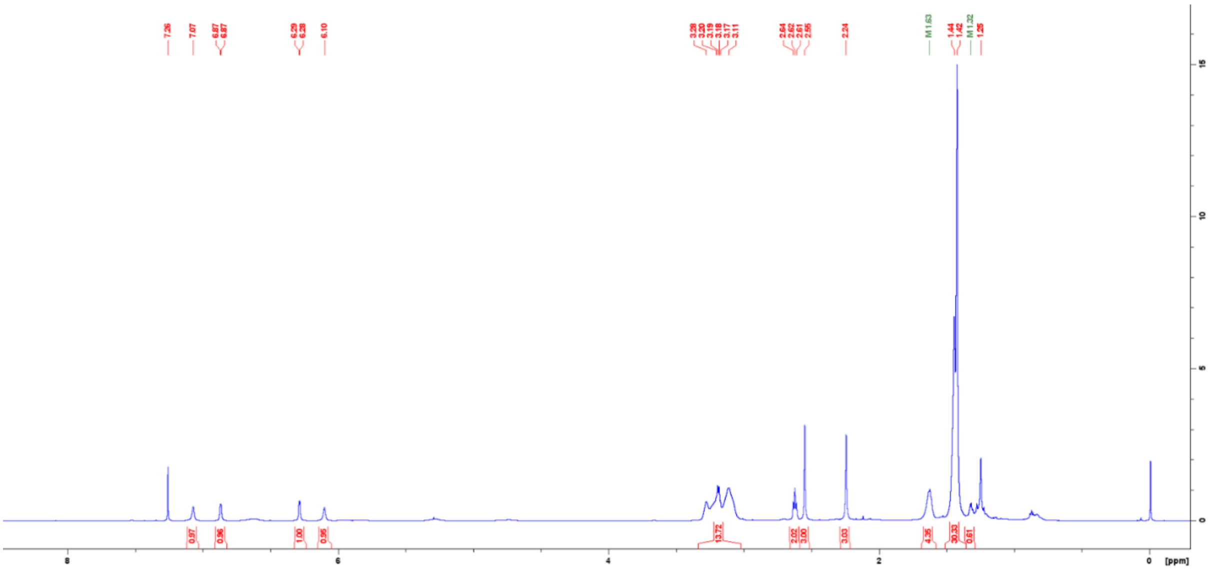

**[13]C-NMR**

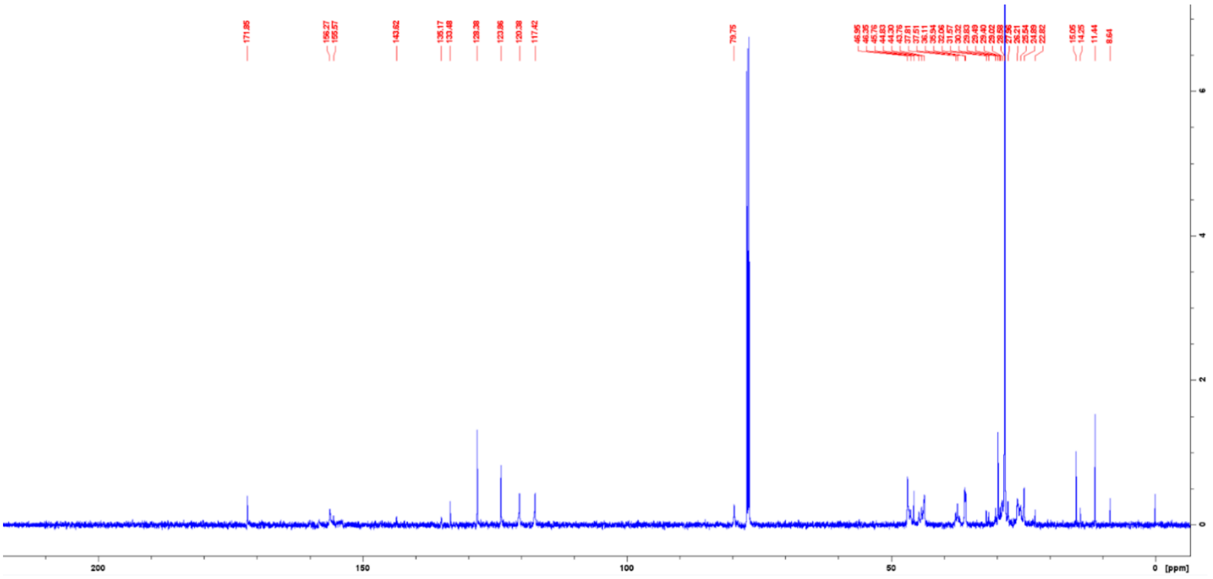

Boc-protected BODIPY-FL-A **7c**

**$^{1}$H-NMR**

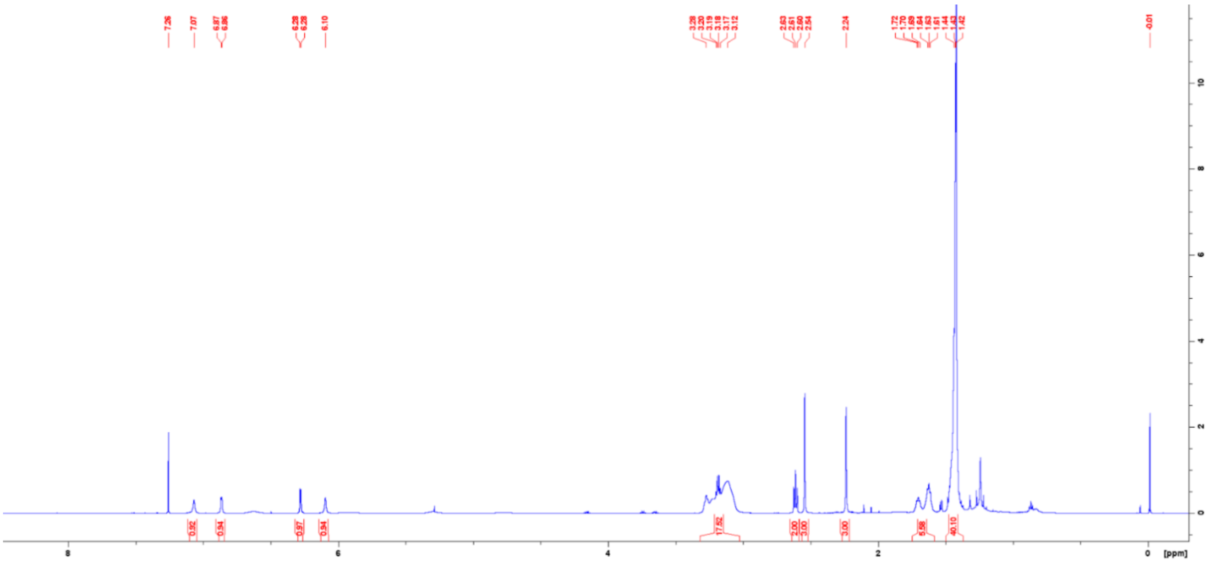

**$^{13}$C-NMR**

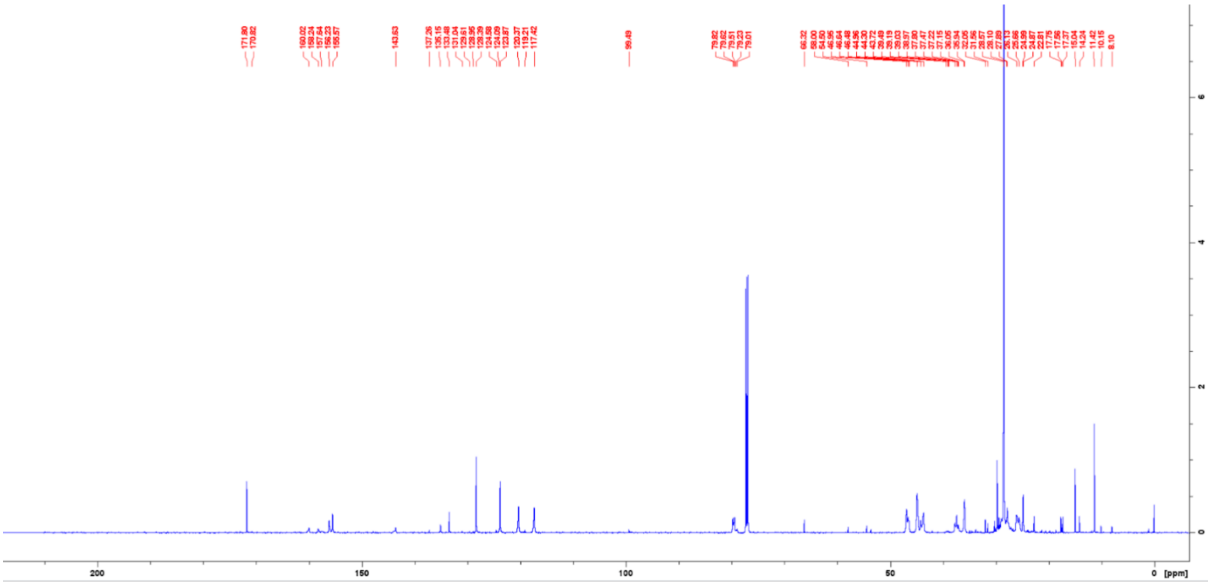

Boc-protected FITC probe **8a**

**[1]H-NMR**

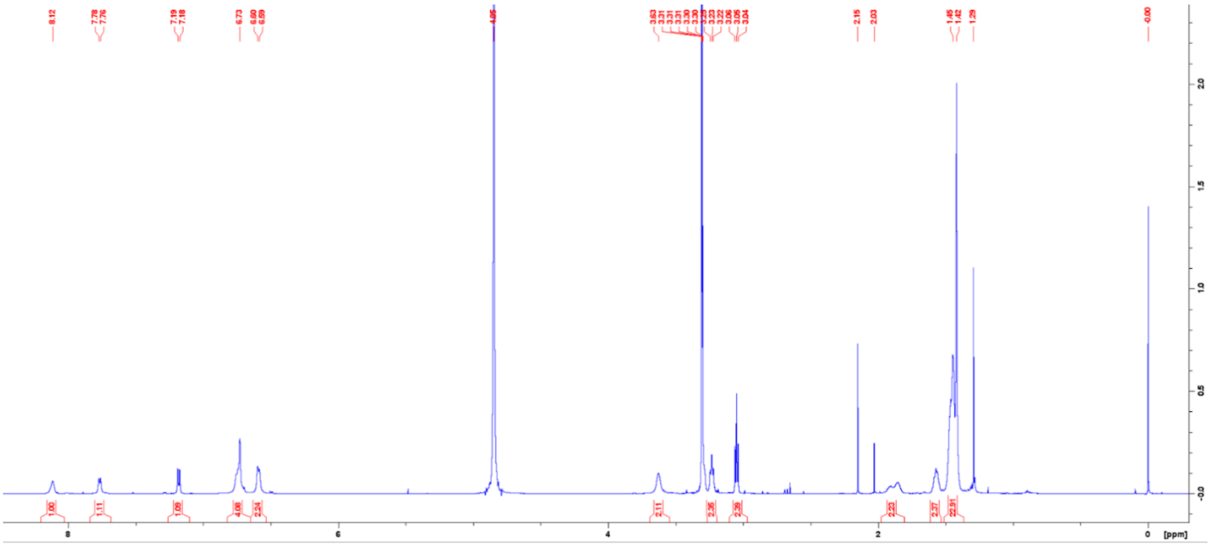

**[13]C-NMR**

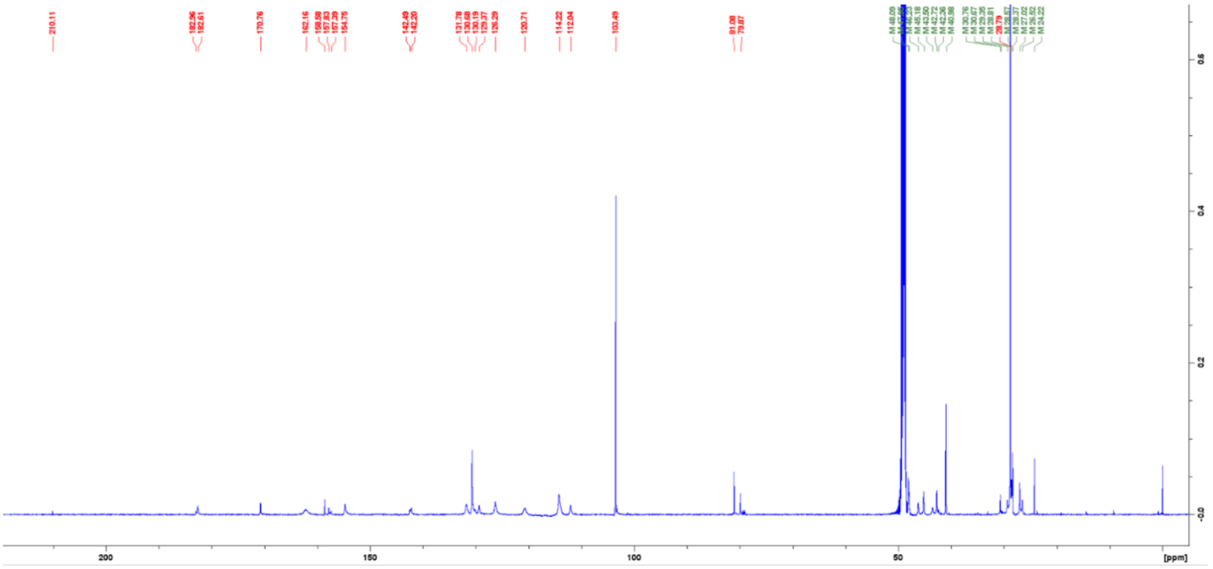

Boc-protected FITC probe **8b**

**$^{1}$H-NMR**

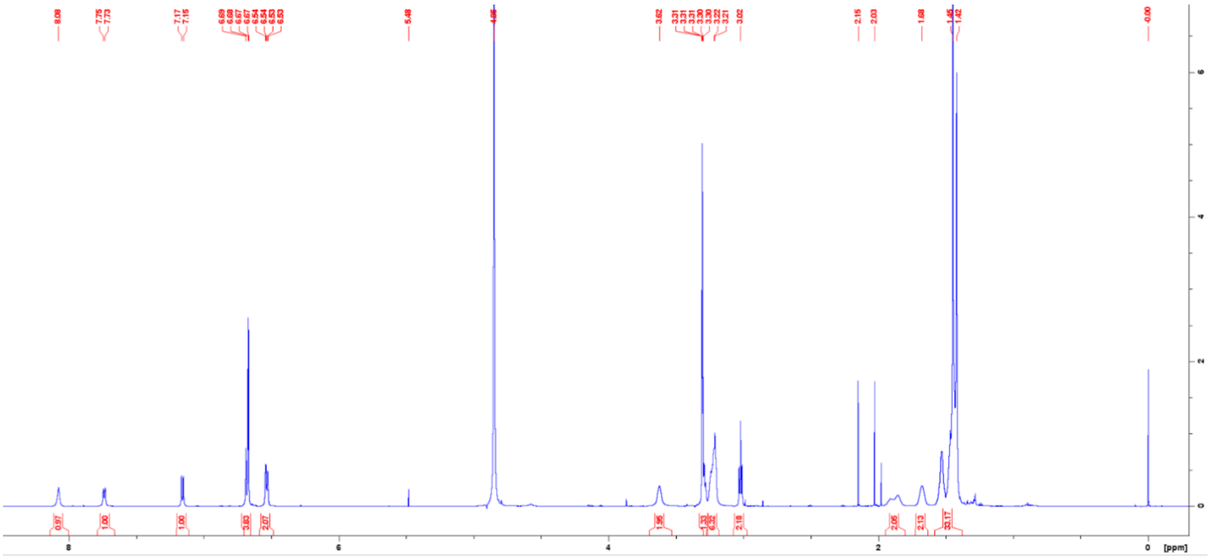

**$^{13}$C-NMR**

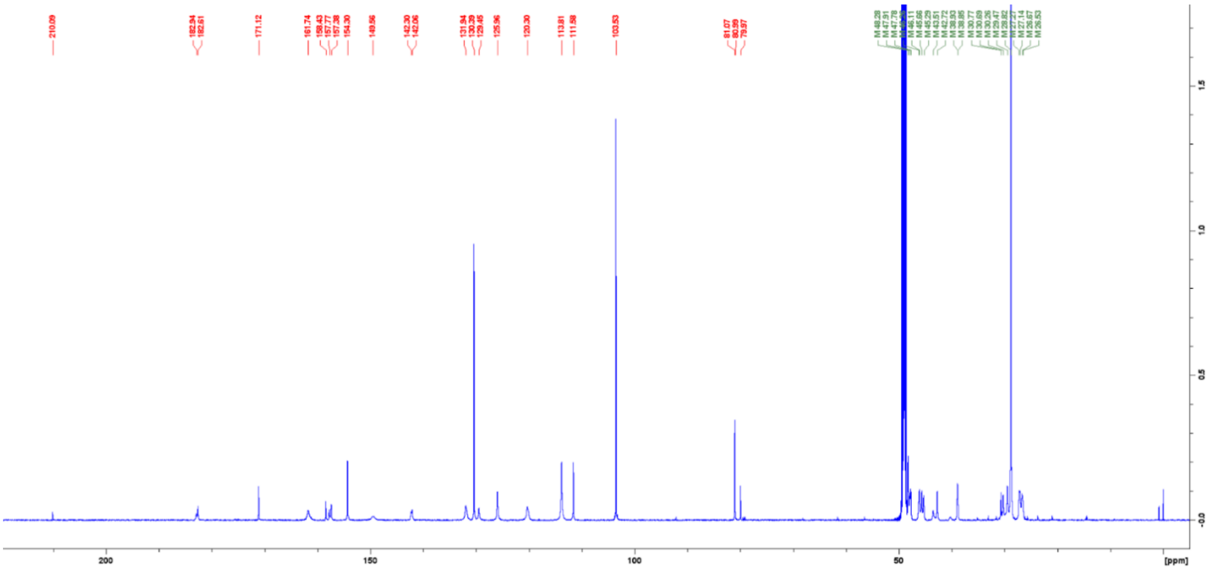

Boc-protected FITC probe **8c**

**$^{1}$H-NMR**

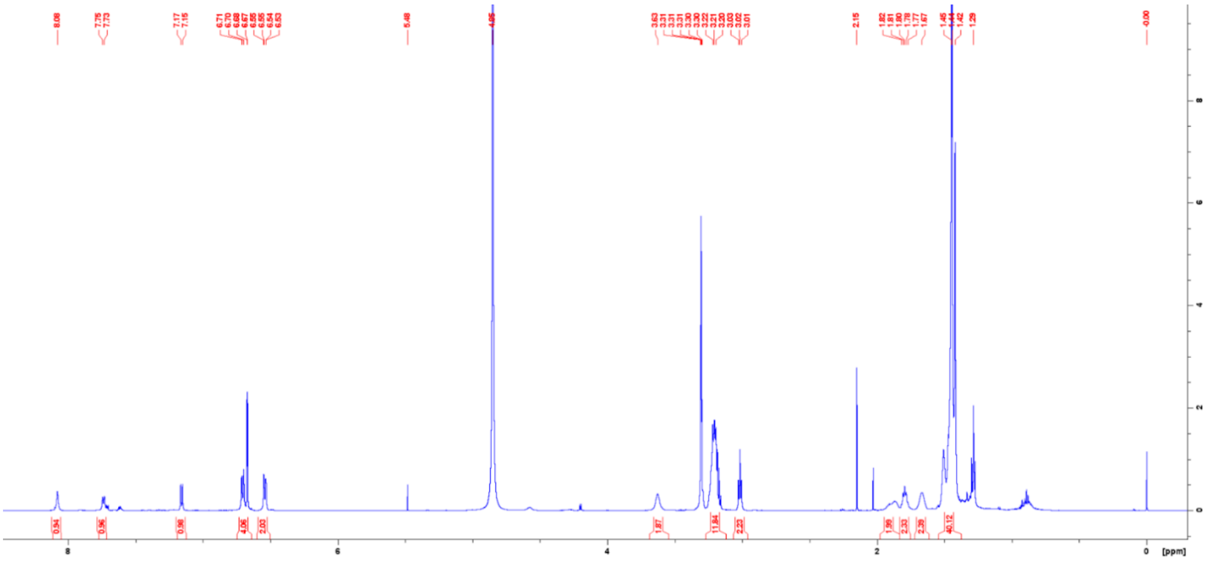


**$^{13}$C-NMR**

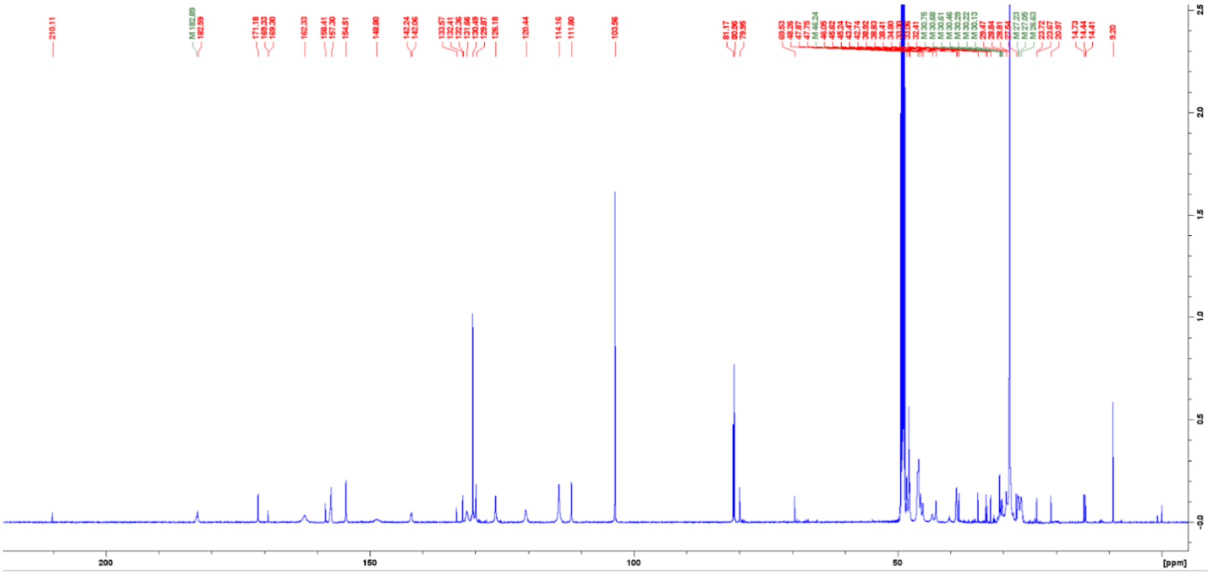

### LC chromatograms and purity of final compounds

BODIPY-FL-T **2a**

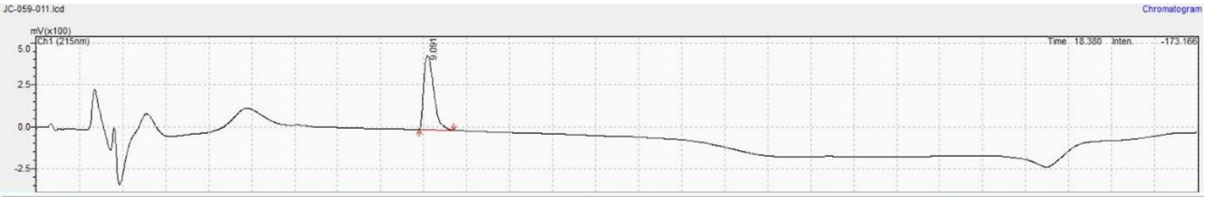


| Peak# | Ret. Time | Area | Height | Mark | Conc. | Unit | ID# | Name | Area% | Conc.% | Norm. Conc. |
|---|---|---|---|---|---|---|---|---|---|---|---|
| 1 | 9.091 | 7094078 | 441546 | M | 0.000 | | | | 100.000 | -- | -- |
| Total | | 7094078 | 441546 | | 0.000 | | | | 100.000 | | |

BODIPY-FL-T **2b**

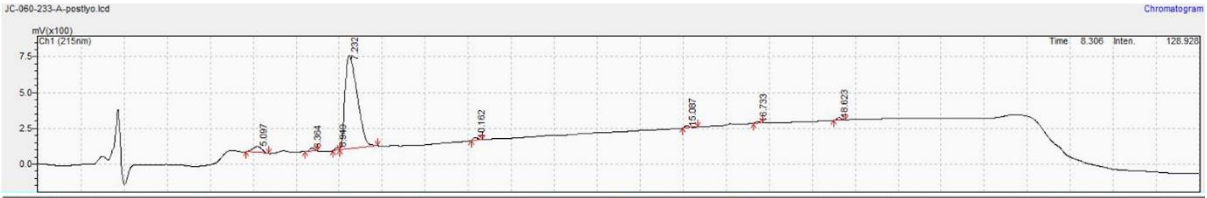


| Peak# | Ret. Time | Area | Height | Mark | Conc. | Unit | ID# | Name | Area% | Conc.% | Norm. Conc. |
|---|---|---|---|---|---|---|---|---|---|---|---|
| 1 | 5.097 | 647528 | 43248 | M | 0.000 | | | | 4.527 | -- | -- |
| 2 | 6.364 | 193155 | 23412 | M | 0.000 | | | | 1.350 | -- | -- |
| 3 | 6.949 | 125130 | 18666 | M | 0.000 | | | | 0.875 | -- | -- |
| 4 | 7.232 | 12935679 | 654721 | V M | 0.000 | | | | 90.428 | -- | -- |
| 5 | 10.162 | 111796 | 15064 | M | 0.000 | | | | 0.782 | -- | -- |
| 6 | 15.087 | 93746 | 11890 | M | 0.000 | | | | 0.655 | -- | -- |
| 7 | 16.733 | 79811 | 11430 | M | 0.000 | | | | 0.558 | -- | -- |
| 8 | 18.623 | 118068 | 16693 | M | 0.000 | | | | 0.825 | -- | -- |
| Total | | 14304913 | 795125 | | 0.000 | | | | 100.000 | | |

BODIPY-FL-T **2c**

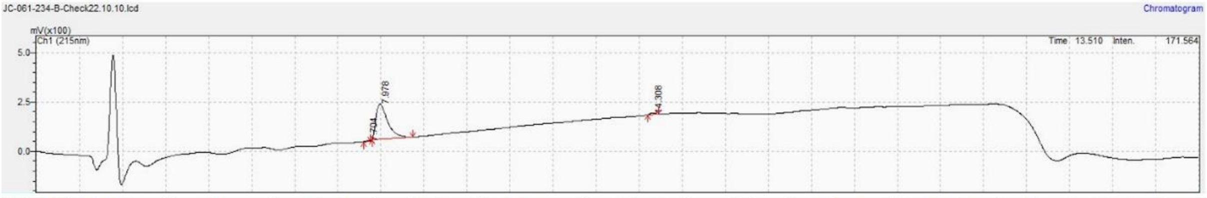


| Peak# | Ret. Time | Area | Height | Mark | Conc. | Unit | ID# | Name | Area% | Conc.% | Norm. Conc. |
|---|---|---|---|---|---|---|---|---|---|---|---|
| 1 | 7.704 | 15292 | 2141 | M | 0.000 | | | | 0.455 | -- | -- |
| 2 | 7.978 | 3297839 | 177394 | M | 0.000 | | | | 98.169 | -- | -- |
| 3 | 14.308 | 46223 | 5359 | M | 0.000 | | | | 1.376 | -- | -- |
| Total | | 3359354 | 184894 | | 0.000 | | | | 100.000 | | |

BODIPY-FL-A **3a**

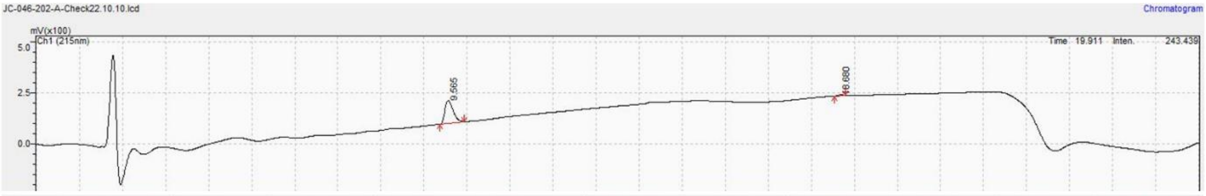


| Peak# | Ret. Time | Area | Height | Mark | Conc. | Unit | ID# | Name | Area% | Conc.% | Norm. Conc. |
|---|---|---|---|---|---|---|---|---|---|---|---|
| 1 | 9.565 | 1473154 | 111020 | M | 0.000 | | | | 97.792 | -- | -- |
| 2 | 18.680 | 33262 | 4437 | M | 0.000 | | | | 2.208 | -- | -- |
| Total | | 1506417 | 115457 | | 0.000 | | | | 100.000 | | |

BODIPY-FL-A **3b**

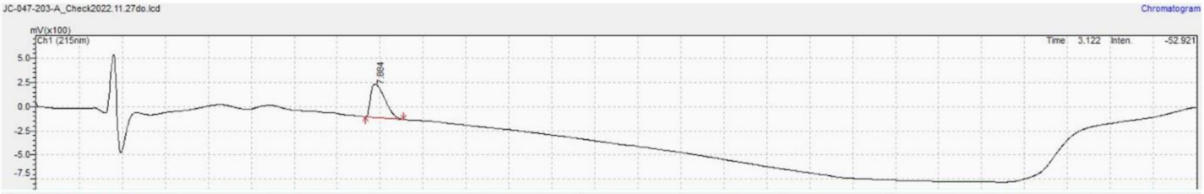


| Peak# | Ret. Time | Area | Height | Mark | Conc. | Unit | ID# | Name | Area% | Conc.% | Norm. Conc. |
|---|---|---|---|---|---|---|---|---|---|---|---|
| 1 | 7.884 | 7877180 | 347052 | M | 0.000 | | | | 100.000 | -- | -- |
| Total | | 7877180 | 347052 | | 0.000 | | | | 100.000 | | |

BODIPY-FL-A **3c**

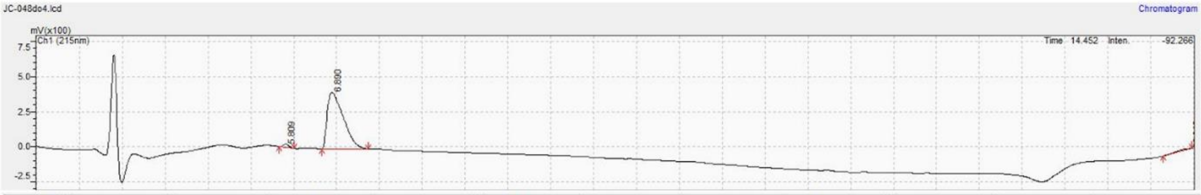


| Peak# | Ret. Time | Area | Height | Mark | Conc. | Unit | ID# | Name | Area% | Conc.% | Norm. Conc. |
|---|---|---|---|---|---|---|---|---|---|---|---|
| 1 | 5.809 | 224126 | 24367 | M | 0.000 | | | | 2.031 | -- | -- |
| 2 | 6.890 | 10567139 | 404039 | M | 0.000 | | | | 95.769 | -- | -- |
| 3 | 26.940 | 242678 | 2195 | | 0.000 | | | | 2.199 | -- | -- |
| Total | | 11033943 | 430601 | | 0.000 | | | | 100.000 | | |

FITC **4a**

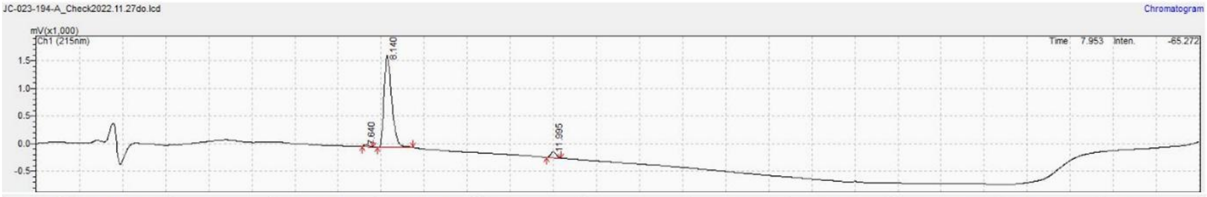


| Peak# | Ret. Time | Area | Height | Mark | Conc. | Unit | ID# | Name | Area% | Conc.% | Norm. Conc. |
|---|---|---|---|---|---|---|---|---|---|---|---|
| 1 | 7.640 | 266878 | 35899 | M | 0.000 | | | | 1.219 | -- | -- |
| 2 | 8.140 | 20730211 | 1668711 | M | 0.000 | | | | 94.699 | -- | -- |
| 3 | 11.995 | 893506 | 103654 | M | 0.000 | | | | 4.082 | -- | -- |
| Total | | 21890595 | 1808264 | | 0.000 | | | | 100.000 | | |

FITC **4b**

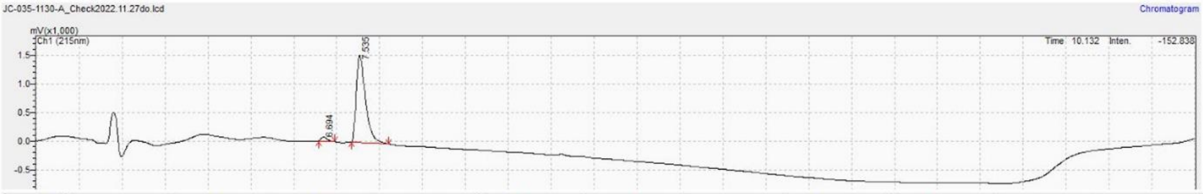


| Peak# | Ret. Time | Area | Height | Mark | Conc. | Unit | ID# | Name | Area% | Conc.% | Norm. Conc. |
|---|---|---|---|---|---|---|---|---|---|---|---|
| 1 | 6.694 | 743759 | 74920 | M | 0.000 | | | | 3.128 | -- | -- |
| 2 | 7.535 | 23035879 | 1533920 | M | 0.000 | | | | 96.872 | -- | -- |
| Total | | 23779638 | 1608840 | | 0.000 | | | | 100.000 | | |

FITC **4c**

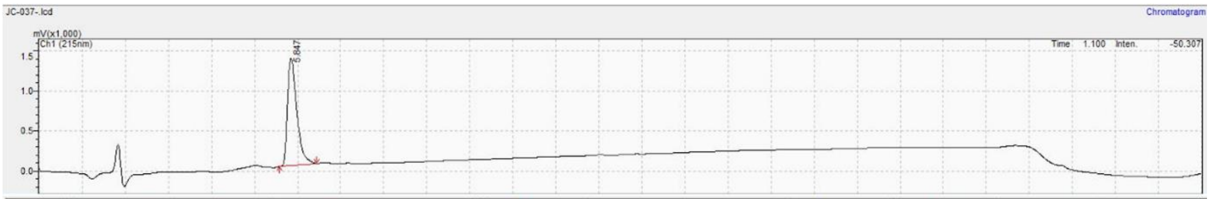


| Peak# | Ret. Time | Area | Height | Mark | Conc. | Unit | ID# | Name | Area% | Conc.% | Norm. Conc. |
|---|---|---|---|---|---|---|---|---|---|---|---|
| 1 | 5.847 | 20326612 | 1340541 | M | 0.000 | | | | 100.000 | -- | -- |
| Total | | 20326612 | 1340541 | | 0.000 | | | | 100.000 | | |

### References Supplementary Information


1. Vanhoutte, R.; Kahler, J.P.; Martin, S.; van Veen, S.; Verhelst, S.H.L. Clickable Polyamine Derivatives as Chemical Probes for the Polyamine Transport System. *Chembiochem* **2018**, *19*, 907-911, doi:10.1002/cbic.201800043.
2. Hamouda, N.N.; Van den Haute, C.; Vanhoutte, R.; Sannerud, R.; Azfar, M.; Mayer, R.; Cortes Calabuig, A.; Swinnen, J.V.; Agostinis, P.; Baekelandt, V.; et al. ATP13A3 is a major component of the enigmatic mammalian polyamine transport system. *J Biol Chem* **2021**, *296*, 100182, doi:10.1074/jbc.RA120.013908.
